# *Culex tarsalis* is a competent host of the insect-specific alphavirus Eilat virus (EILV)

**DOI:** 10.1101/2022.08.06.503036

**Authors:** Renuka E. Joseph, Nadya Urakova, Kristine L. Werling, Hillery C. Metz, Kaylee Montanari, Jason L. Rasgon

## Abstract

Eilat virus (EILV) is an insect-specific alphavirus that has the potential to be developed into a tool to combat mosquito-borne pathogens. However, its mosquito host range and transmission routes are not well understood. Here we fill this gap by investigating EILV’s host competence and tissue tropism in five mosquito species: *Aedes aegypti, Culex tarsalis, Anopheles gambiae, Anopheles stephensi*, and *Anopheles albimanus*. Of the tested species, *Cx. tarsalis was* the most competent host for EILV. The virus was found in *Cx. tarsalis* ovaries, but no vertical or venereal transmission was observed. *Culex tarsalis* also transmitted EILV via saliva, suggesting the potential for horizontal transmission between an unknown vertebrate or invertebrate host. We found that reptile (turtle and snake) cells lines were not competent for EILV infection. We tested a potential invertebrate host (*Manduca sexta* caterpillars) but found they were not susceptible to EILV infection. Together, our results suggest that EILV could be developed as a tool to target pathogenic viruses that use *Culex tarsalis* as a vector. Our work sheds light on the infection and transmission dynamics of a poorly understood insect-specific virus and reveals it may infect a broader range of mosquito species than previously recognized.

**IMPORTANCE:** The recent discovery of insect-specific alphaviruses presents opportunities both to study the biology of virus host range and to develop them into tools against pathogenic arboviruses. Here we characterize the host range and transmission of Eilat virus in five mosquito species. We find that *Culex tarsalis*—a vector of harmful human pathogens including West Nile Virus—is a competent host of Eilat virus. However, how this virus is transmitted between mosquitoes remains unclear. We find that Eilat virus infects the tissues necessary for both vertical and horizontal transmission—a crucial step in discerning how Eilat virus maintains itself in nature.

## INTRODUCTION

The small, spherical, enveloped positive-sense RNA viruses in the genus *Alphavirus* (family *Togaviridae*) are primarily mosquito-borne and include important human pathogens such as Mayaro, O’nyong-nyong, Chikungunya, and Ross River viruses^1^. The 11-12 kb alphavirus RNA genome has two open reading frames (ORFs): The 5’ end of the genome encodes four non-structural proteins (nsP1-4), and the *3*’ end that encodes five structural proteins (sPs; Capsid, E3, E2, 6K, and E1) expressed by a sub-genomic promoter^1^. Alphaviruses typically have a broad host range spanning vertebrates such as humans, non-human primates, horses, birds, reptiles, amphibians, as well as invertebrates such as mosquitoes, ticks, and lice^2^. Horizontal transmission between mosquitoes and vertebrates is how alphaviruses typically maintain themselves in nature^2^. However, several insect-specific alphaviruses that cannot infect vertebrate cells have been recently discovered^3–6^. Though their host range and transmission route(s) remain poorly described, a better understanding of these host-restricted viruses may lead to new insights into virus biology, and ultimately to the development of new tools for curbing the spread of mosquito-borne pathogens.

Eilat virus (EILV) is an insect-specific alphavirus originally isolated from *Anopheles coustani* mosquitoes collected during an arbovirus survey in the Negev desert of Israel^3^. Phylogenetically, EILV clusters with the mosquito-borne clade of the *Alphavirus* genus, basal to the western equine encephalitis complex^3^. Nasar *et al.* were the first to characterize EILV and found that it infects and replicates in the insect cell lines C6/36 and C7/10 (*Aedes albopictus*), CT (*Culex tarsalis*), and PP-9 (*Phlebotomus papatasi*), but not in mammalian, avian or amphibian cell lines^3^—indicating it is an insect specific virus (ISV). Similarly, EILV could not infect newborn mice, a model for alphavirus infections^7^. EILV host range restriction was found to occur at both the attachment/entry and the viral genome replication levels in vertebrates^7^.

A broad range of mosquito species act as vectors for different alphaviruses including members of the genera *Aedes, Anopheles*, and *Culex*^2,8^. In previous work, four mosquito species were found to be susceptible to EILV infection to varying degrees^9^. *Ae. aegypti* was the most susceptible following oral challenge, while *Ae. albopictus, An. gambiae*, and *Cx. quinquefasciatus* were only susceptible to EILV at the highest dose tested (10^9^ PFU/ml)^8^. EILV post-oral infections in *Ae. aegypti, An. gambiae* and *Cx. quinquefasciatus* (but not Ae. *albopictus*) were also able to disseminate beyond the midgut^8^. Nevertheless, these findings suggested that EILV has a restricted host range, as all but one of the examined species (Ae. *aegypti*) were refractory to infection at titers typical for other alphaviruses^9,10^.

ISVs such as *Culex* flaviviruses (CxFV) are thought to be adapted to a single host system, within which they are transmitted vertically from mothers to offspring^11,12^. Vertical transmission routes used by ISVs include transovarial (viral infection of germline tissue in mosquito ovaries) and transovular transmission (viral infection of mosquito eggs as they pass the oviduct)^11–13^. In contrast, horizontal transmission can occur when a virus infects the mosquito salivary glands and is subsequently passed to a new host by salivation during feeding^2^. Thus, knowledge of tissue tropism can shed light on viral transmission routes. In the case of EILV, Nasar *et al.* found that that midguts of four mosquito species were infected following intrathoracic injection with the virus^8^. EILV was found in the salivary glands of *Ae. aegypti, An. gambiae*, and *Cx. quinquefasciatus* but was not detected in the ovaries of the tested species, suggesting the potential for horizontal but not vertical transmission.

The absence of EILV in the ovaries is unexpected for an insect-specific virus (ISV) as it calls into question how this virus is transmitted between mosquitoes. However, EILV was detected in both adult and larval *Culex pipiens* in Morocco, consistent with the hypothesis that EILV uses transovarial or transovular transmission routes^14^, typical of an ISV, though environmental transmission is an alternative explanation. The failure to detect EILV in mosquito ovaries in a laboratory setting thus far indicates a need for more research into its transmission route(s).

Alternately, the presence of EILV in the salivary glands suggests horizontal transmission to an unknown host such as an uncharacterized vertebrate or perhaps another invertebrate. Reptiles have not yet been assessed for their competence to host EILV. Additionally, though it is not a well-characterized transmission route, George *et al.* showed that *Anopheles stephensi* are attracted to and can successfully feed on the caterpillars *Manduca sexta* and *Heliothis subflexa*^15^. This raises the possibility that these caterpillars may play a role in the transmission of EILV or other ISVs. To address these questions and gaps, here we investigate i) the ability of *Aedes aegypti, Culex tarsalis, Anopheles gambiae, Anopheles stephensi*, and *Anopheles albimanus* to become infected, disseminate, and transmit EILV, ii) the tissue tropism and transmission route of EILV in these species and iii) the susceptibility of reptile cell lines and *Manduca sexta* to EILV infection.

## RESULTS

### *Culex tarsalis* is a competent host of orally acquired EILV

To determine host competence of each species, 102 *Ae. aegypti* (Rockefeller strain), 78 *Cx. tarsalis* (YOLO strain), 100 *An. gambiae* (Keele strain), 100 *An. stephensi* (Liston strain), and *83 An. albimanus* (STECLA strain) mosquitoes were orally challenged with EILV-eGFP (Table 1). The infection rate (IR) was defined as the proportion of infected mosquitoes among the total number of engorged mosquitoes, the dissemination rate (DIR) the proportion of infected mosquitoes with virus-positive legs, the transmission rate (TR) the proportion of mosquitoes with virus-positive saliva among those with virus-positive legs, and the transmission efficiency (TE) the proportion of mosquitoes with virus-positive saliva among the total number of mosquitoes engorged.

**TABLE 1.**
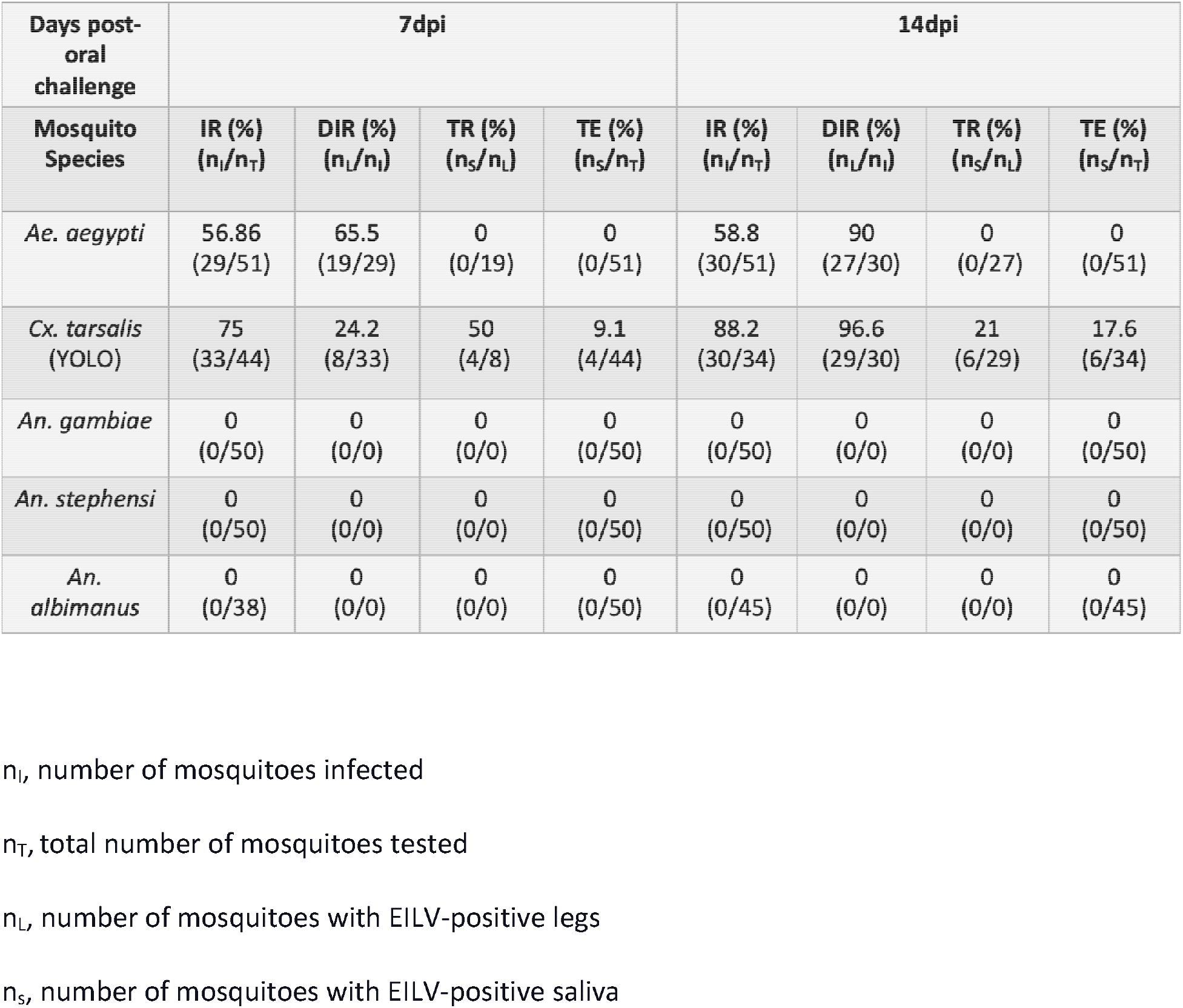
The infection (IR), dissemination (DIR) and transmission (TR) rates, and transmission efficiency (TE) of five mosquito species orally challenged with EILV-eGFP

| Days post-oral challenge | 7dpi |  |  |  | 14dpi |  |  |  |
| --- | --- | --- | --- | --- | --- | --- | --- | --- |
| Mosquito Species | IR (%)<br>($n_I/n_T$) | DIR (%)<br>($n_L/n_I$) | TR (%)<br>($n_S/n_L$) | TE (%)<br>($n_S/n_T$) | IR (%)<br>($n_I/n_T$) | DIR (%)<br>($n_L/n_I$) | TR (%)<br>($n_S/n_L$) | TE (%)<br>($n_S/n_T$) |
| *Ae. aegypti* | 56.86<br>(29/51) | 65.5<br>(19/29) | 0<br>(0/19) | 0<br>(0/51) | 58.8<br>(30/51) | 90<br>(27/30) | 0<br>(0/27) | 0<br>(0/51) |
| *Cx. tarsalis*<br>(YOLO) | 75<br>(33/44) | 24.2<br>(8/33) | 50<br>(4/8) | 9.1<br>(4/44) | 88.2<br>(30/34) | 96.6<br>(29/30) | 21<br>(6/29) | 17.6<br>(6/34) |
| *An. gambiae* | 0<br>(0/50) | 0<br>(0/0) | 0<br>(0/0) | 0<br>(0/50) | 0<br>(0/50) | 0<br>(0/0) | 0<br>(0/0) | 0<br>(0/50) |
| *An. stephensi* | 0<br>(0/50) | 0<br>(0/0) | 0<br>(0/0) | 0<br>(0/50) | 0<br>(0/50) | 0<br>(0/0) | 0<br>(0/0) | 0<br>(0/50) |
| *An. albimanus* | 0<br>(0/38) | 0<br>(0/0) | 0<br>(0/0) | 0<br>(0/50) | 0<br>(0/45) | 0<br>(0/0) | 0<br>(0/0) | 0<br>(0/45) |

$n_I$ , number of mosquitoes infected
$n_T$ , total number of mosquitoes tested
$n_L$ , number of mosquitoes with EILV-positive legs
$n_S$ , number of mosquitoes with EILV-positive saliva

We found that EILV infected two species, *Ae. aegypti* and *Cx. tarsalis*, with infection rates (IRs) in the range of 57-88% at both 7 and 14 days post infection (dpi). In contrast, *An. gambiae, An. stephensi, and An. albimanus* were refractory to oral infection with EILV. Of the two susceptible species, *Cx. tarsalis* was more likely to become infected. Specifically, the IR of *Cx. tarsalis* at 14 dpi was significantly greater than that of Ae. *aegypti* (Fisher’s exact test, P≤0.01), though at 7 dpi the species did not differ (Fisher’s exact test, P>0.05). Within each species, IRs did not significantly change over time (i.e., 7 dpi *versus* 14 dpi; Table 1; Fishers exact tests, P>0.05 for both).

EILV infections disseminated beyond the midgut at both time points in both infected species, but the dissemination rate (DIR) of Ae. *aegypti* was significantly higher (Fisher’s exact test, P≤0.01) than that of *Cx. tarsalis* at 7 dpi. However, this difference disappeared by 14 dpi (P>0.05) due to increased dissemination in *Cx. tarsalis*. Only *Cx. tarsalis* had EILV positive saliva, with the transmission efficiency (TE) rising over time from 9.1% (7 dpi) to 17.6% (14 dpi), while the transmission rate (TR) dropped over time from 50% to 21% due to the increase in number of disseminated infections at the later time point (Table 1). The mean EILV titers in positive saliva samples at 7 dpi and 14 dpi were 5.82 and 4.58 FFU/mosquito, respectively.

At the level of titer, *Ae. aegypti* and *Cx. tarsalis* (YOLO) body and leg samples did not differ at 7 dpi (Fig. 1; Mann-Whitney U tests, P>0.05 for both comparisons). However, by 14 dpi the EILV titers in *Cx. tarsalis* body and leg samples were significantly greater (Fig. 1; P≤0.0001 for both) than those of Ae. *aegypti*. Together, our results show that *Cx. tarsalis* is a competent transmitting host for EILV via oral route of infection.

**FIG 1.**
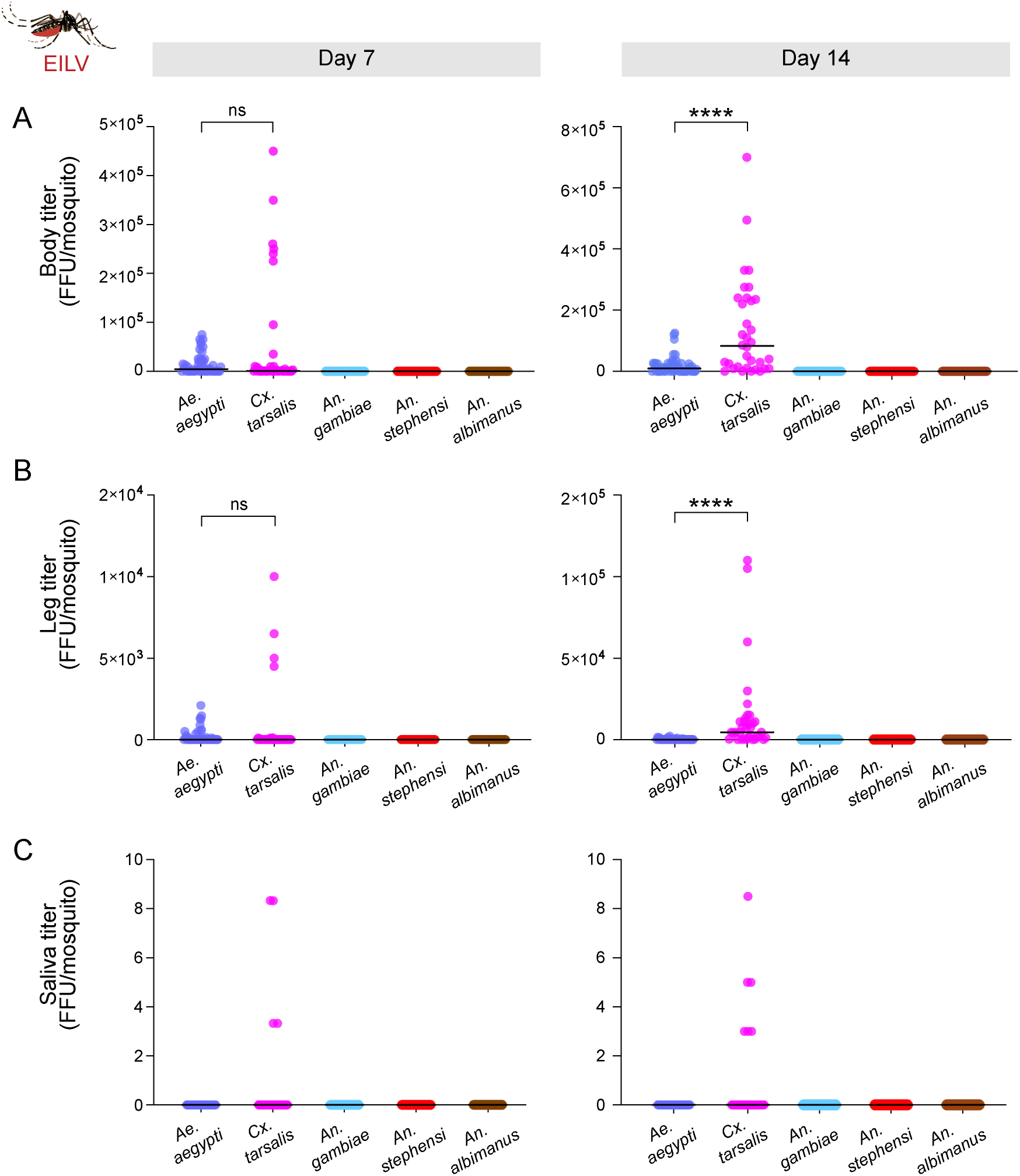
Viral titers of body, leg, and saliva samples from five mosquito species orally challenged with infectious EILV-eGFP. Titers are plotted at 7 and 14 dpi for (A) body, (B) leg and (C) saliva samples collected from mosquitoes orally challenged with EILV-eGFP (10^7^ FFU/ml). Each point represents a single mosquito sample, while horizontal bars depict the group medians. Significance was evaluated using Mann-Whitney U tests. **** P<0.0001.

### Genetically diverse *Cx. tarsalis* strains are susceptible to EILV infection and may transmit it via saliva

Having established that a lab strain of *Cx. tarsalis* (YOLO) is a competent host for EILV, we next asked if this susceptibility is strain specific (i.e., limited to YOLO), or rather, widespread across diverse colonies of this species. We therefore assessed the ability of EILV to infect an additional lab colony of this species (KNWR strain), as well as *Cx. tarsalis* recently captured from the wild in California.

We orally challenged 73 KNWR and 63 wild *Cx. tarsalis* with EILV-eGFP and tested body, leg, and saliva samples for EILV at 7 and 14 dpi to determine the IR, DIR, TR, and TE of both strains and compare these values to those of YOLO (Table 2). We found the KNWR colony had similar susceptibility to EILV infection as YOLO. Specifically, their IRs did not differ at 7 or 14 dpi (Fisher’s exact tests, P>0.05 for both). In contrast, wild *Cx. tarsalis* were less susceptible to EILV infection, with IRs significantly lower than those YOLO across time (Fisher’s exact tests, 7 dpi and 14 dpi, both P≤0.0001). We found differences in dissemination were limited to late infections—no differences in DIR were found at 7 dpi (P>0.05 for both strain comparisons), but by 14 dpi, both KNWR and wild *Cx. tarsalis* had lower rates than YOLO (both P≤0.05).

**TABLE 2.** The infection (IR), dissemination (DIR) and transmission (TR) rates and transmission efficiency (TE) of *Cx. tarsalis* mosquitoes orally challenged with EILV-eGFP

| Days post-oral challenge | 7dpi |  |  |  | 14dpi |  |  |  |
| --- | --- | --- | --- | --- | --- | --- | --- | --- |
| *Cx. tarsalis* strain | IR (%)<br>($n_i/n_T$) | DIR (%)<br>($n_L/n_i$) | TR (%)<br>($n_s/n_L$) | TE (%)<br>($n_s/n_T$) | IR (%)<br>($n_i/n_T$) | DIR (%)<br>($n_L/n_i$) | TR (%)<br>($n_s/n_L$) | TE (%)<br>($n_s/n_T$) |
| *Cx. tarsalis* (YOLO) | 75<br>(33/44) | 24.2<br>(8/33) | 50<br>(4/8) | 9.1<br>(4/44) | 88.2<br>(30/34) | 96.6<br>(29/30) | 21<br>(6/29) | 17.6<br>(6/34) |
| *Cx. tarsalis* (KNWR) | 76.3%<br>(29/38) | 27.5<br>(8/29) | 37.5<br>(3/8) | 7.9<br>(3/38) | 68.6<br>(24/35) | 71<br>(17/24) | 35.3<br>(6/17) | 17.1<br>(6/35) |
| Wild *Cx. tarsalis* | 22<br>(7/31) | 57<br>(4/7) | 25<br>(1/4) | 3.2<br>(1/31) | 25<br>(8/32) | 62.5<br>(5/8) | 0<br>(0/5) | 0<br>(0/32) |

$n_i$ , number of mosquitoes infected
$n_T$ , total number of mosquitoes tested
$n_L$ , number of mosquitoes with EILV-positive legs
$n_s$ , number of mosquitoes with EILV-positive saliva

EILV was found in the saliva of all assayed *Cx. tarsalis* strains. The TR and TE of KNWR and YOLO at 7 dpi and 14 dpi did not differ (Fisher’s exact tests, all P>0.05). Similarly, the TR and TE of wild *Cx. tarsalis* and YOLO were statistically indistinguishable at 7 dpi (both P>0.05), though wild mosquitoes had a lower TE (but not TR) at 14 dpi compared to YOLO (P≤0.05).

There were additional differences in EILV host competence at the viral titer level. The titers in the body and leg samples of KNWR did not differ from those of YOLO at 7 dpi (Mann-Whitney U test, both P>0.05), but were lower at 14 dpi (P≤0.01 for both) (Fig. 2A-B). The leg and body titers of wild *Cx. tarsalis were* significantly lower than those of YOLO in most comparisons (see Fig. 2 for full results). EILV titers in saliva samples did not differ between any *Cx. tarsalis* strains at either time point (all P>0.05), though EILV was absent in the saliva of wild *Cx. tarsalis* at 14 dpi. Together, these results demonstrate that genetically diverse *Cx. tarsalis* mosquitoes are susceptible to EILV via oral route of infection, though there was variation among strains with regards to EILV infection dynamics—with *Cx. tarsalis* (YOLO) being the most susceptible and wild *Cx. tarsalis* the least susceptible to EILV infection.

**FIG 2.**
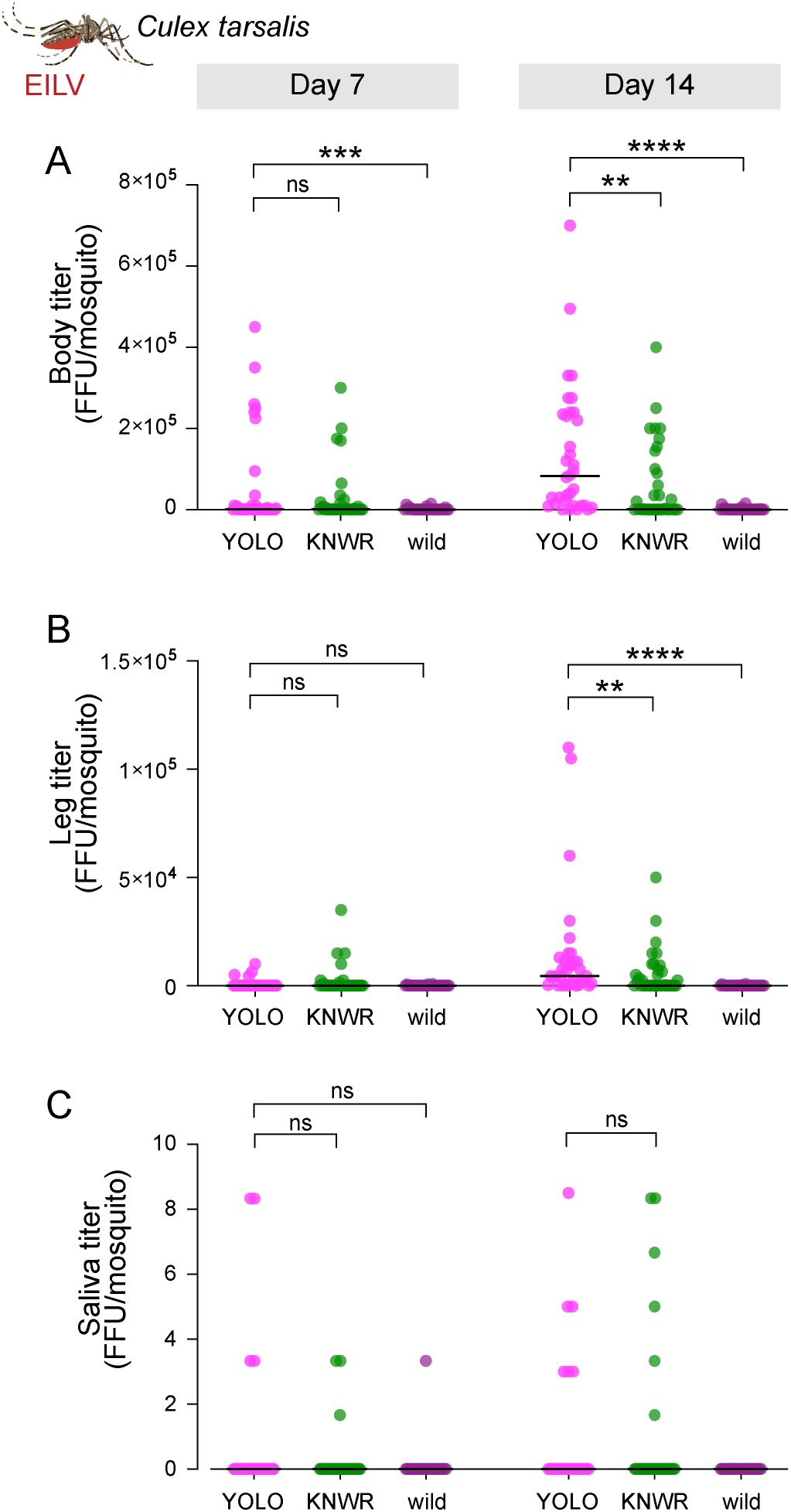
Viral titers of body, leg, and saliva samples from three *Cx. tarsalis* strains orally challenged with EILV-eGFP. Titers are plotted at 7 and 14 dpi for (A) body, (B) leg and (C) saliva samples from mosquitoes orally challenged with EILV-eGFP (10^7^ FFU/ml). Each point represents a single mosquito sample, while horizontal bars depict the group medians. Note that data for *Cx. tarsalis* (YOLO) is re-plotted from Fig. 1. Statistical significance was evaluated using Mann-Whitney U tests. ** P<0.01, *** P<0.001, **** P<0.0001.

### Midgut infection barriers block EILV infections in *Anopheles*

By injecting virus directly into the thorax, we next asked if EILV host competence is broadened when any tissue-specific barriers to infection (e.g., midgut barriers) are bypassed. We therefore examined EILV presence in bodies at 7 and 14 dpi following intrathoracic (IT) injections in 74 *Ae. aegypti*, 80 *Cx. tarsalis* (YOLO), *66 An. gambiae, 76 An. stephensi*, and *73 An. albimanus* (Table 3). Whereas only two species were susceptible to oral infections, all five mosquito species were susceptible to EILV infection when challenged intrathoracically, implying the existence of a strong midgut infection barrier to EILV infection in the tested anophelines. Following IT injections, the IR for *Ae. aegypti, Cx*. tarsalis, and *An. gambiae* was 100% at both time points, while *An. stephensi* had a slightly lower rate of 92.7% and 94.3% at 7 and 14 dpi, respectively (Table 3). *An. albimanus* had the lowest infection rates—94.7% and 85.7% at 7 and 14 dpi, respectively (Table 3). EILV IR did not differ by species or time point (Fisher’s exact tests, all P>0.05). However, EILV titers of *Cx. tarsalis* (YOLO) body samples at 7 and 14 dpi were significantly higher than those of the other tested species (Fig. 3A), consistent with this species being the most vulnerable to EILV infection.

**FIG 3.**
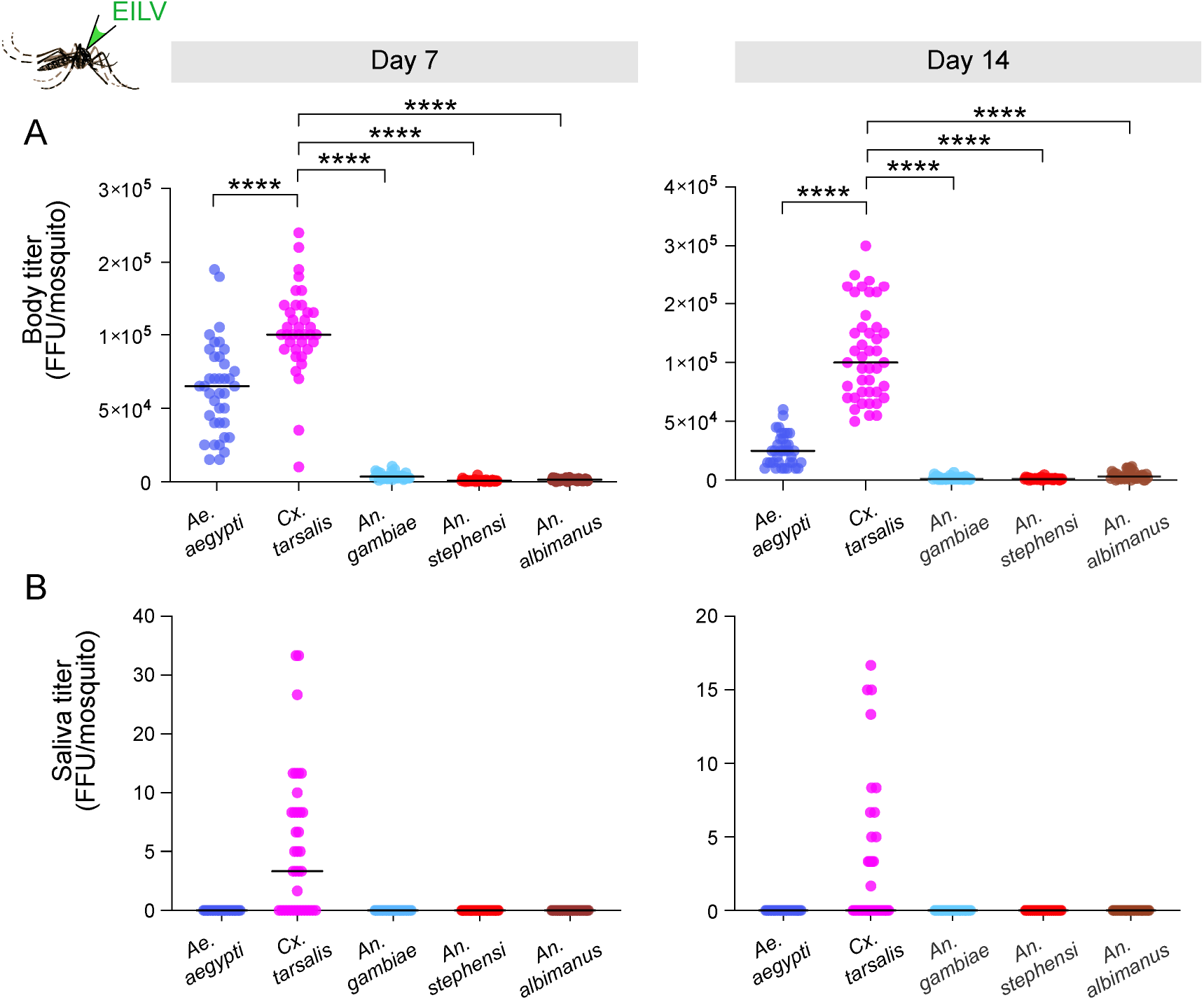
Viral titers of body, leg, and saliva samples from five mosquito species intrathoracically injected with EILV-eGFP. Titers are plotted at 7 and 14 dpi for (A) body and (B) saliva samples following IT injection with EILV-eGFP (10^7^ FFU/ml). Each point represents a single mosquito sample and horizontal bars depict group medians. Statistical significance was evaluated using Mann-Whitney U test. **** P<0.0001.

**TABLE 3.** The infection rates (IR) and transmission efficiency (TE) of intrathoracically injected mosquitoes

| Days post-IT challenge | 7dpi |  | 14dpi |  |
| --- | --- | --- | --- | --- |
| Mosquito Species | IR (%)<br>($n_I/n_T$) | TE (%)<br>($n_S/n_T$) | IR (%)<br>($n_I/n_T$) | TE (%)<br>($n_S/n_T$) |
| *Ae. aegypti* | 100<br>(38/38) | 0<br>(0/38) | 100<br>(36/36) | 0<br>(0/36) |
| *Cx. tarsalis*<br>(YOLO) | 100<br>(37/37) | 62.2<br>(23/37) | 100<br>(43/43) | 35<br>(23/43) |
| *An. gambiae* | 100<br>(33/33) | 0<br>(0/33) | 100<br>(33/33) | 0<br>(0/33) |
| *An. stephensi* | 92.7<br>(38/41) | 0<br>(0/41) | 94.3<br>(33/35) | 0<br>(0/35) |
| *An. albimanus* | 94.7<br>(36/38) | 0<br>(0/38) | 85.7<br>(30/35) | 0<br>(0/35) |

$n_I$ , number of mosquitoes infected
$n_T$ , total number of mosquitoes tested
$n_S$ , number of mosquitoes with EILV-positive saliva

### Tissue-specific barriers block EILV from infecting salivary glands in *Aedes* and *Anopheles*

By bypassing the midgut barriers to infection by injecting virus directly into the thorax, EILV had access to the salivary glands of all tested mosquito species, allowing us to test whether EILV can enter their saliva. We examined saliva samples from the five species following intrathoracic (IT) injections for the presence of EILV. As with oral infections, EILV was only found in the saliva of *Cx. tarsalis* following IT challenge, where it was prevalent at both 7 and 14 dpi (Fig. 3B and Table 3; TE of 62.2% and 35% at 7 and 14 dpi). The mean titers of virus positive saliva samples at 7 and 14 dpi were 7.3 and 6.0 FFU/mosquito, respectively (Fig. 3B). Overall, our results show that *Cx. tarsalis* has limited barriers to EILV infection compared to the other mosquito species in our study and is the most competent host for EILV post IT infection.

### EILV is present in the ovaries and salivary glands of orally infected *Cx. tarsalis* mosquitoes

Following an infectious bloodmeal, we used fluorescence microscopy to examine the tissue tropism of EILV-eGFP across the body of all five species at 7 and 14 dpi. All examined tissues (i.e., midgut, salivary glands, and ovaries) of orally challenged *An. gambiae, An. albimanus* and *An. stephensi* were negative for eGFP expression (Fig. 4). However, eGFP expression was detected in the midgut and salivary glands of all three *Cx. tarsalis* strains (YOLO, KNWR and wild) and in *Ae. aegypti* at 7 and 14 dpi. Expression of eGFP in ovary was only observed in two of the *Cx. tarsalis* strains (YOLO and KNWR) and was not observed in wild *Cx. tarsalis* or Ae. *aegypti* at either time point (Fig. 4).

**FIG 4.**
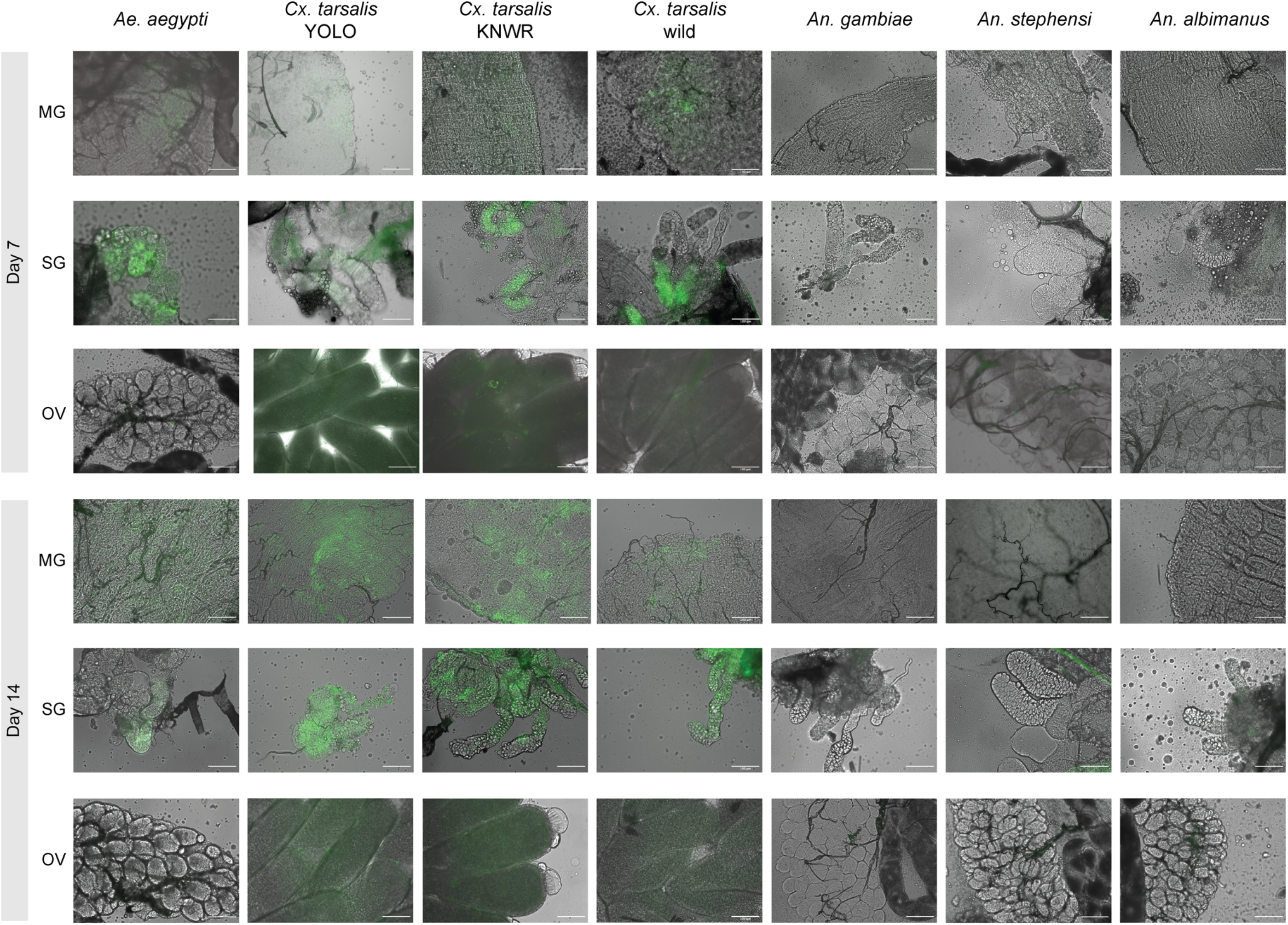
EILV tissue tropism at 7 and 14 dpi in orally challenged mosquitoes. Representative images show eGFP fluorescence (or its absence) in midguts (MG), salivary glands (SG) and ovaries (OV) of mosquitoes fed an infectious bloodmeal containing EILV-eGFP (10^7^ FFU/ml). The brightfield and FITC images have been merged. All scale bars equal 100 μm.

### EILV is absent in the ovaries of *Cx. tarsalis* but present in the ovaries *of An. gambiae* post IT injection

We additionally examined EILV tissue tropism in mosquitoes injected with the virus. Strong eGFP expression—indicating EILV infection—was detected in the midguts of IT injected *Cx. tarsalis* (YOLO) and *Ae. aegypti* at both 7 and 14 dpi (Fig. 5). Similarly, IT injected *An. gambiae, An. stephensi* and *An. albimanus* midguts showed limited eGFP expression at 7 and 14 dpi (Fig. 5). eGFP was also observed in the salivary glands of *Cx. tarsalis* and *Ae. aegypti* but not in the salivary glands of any of the anophelines at either time point (Fig. 5). Surprisingly, the ovaries of IT injected *An. gambiae* were positive for EILV, but the ovaries of the other four species were negative for EILV at 7 and 14 dpi (Fig. 5).

**FIG 5.**
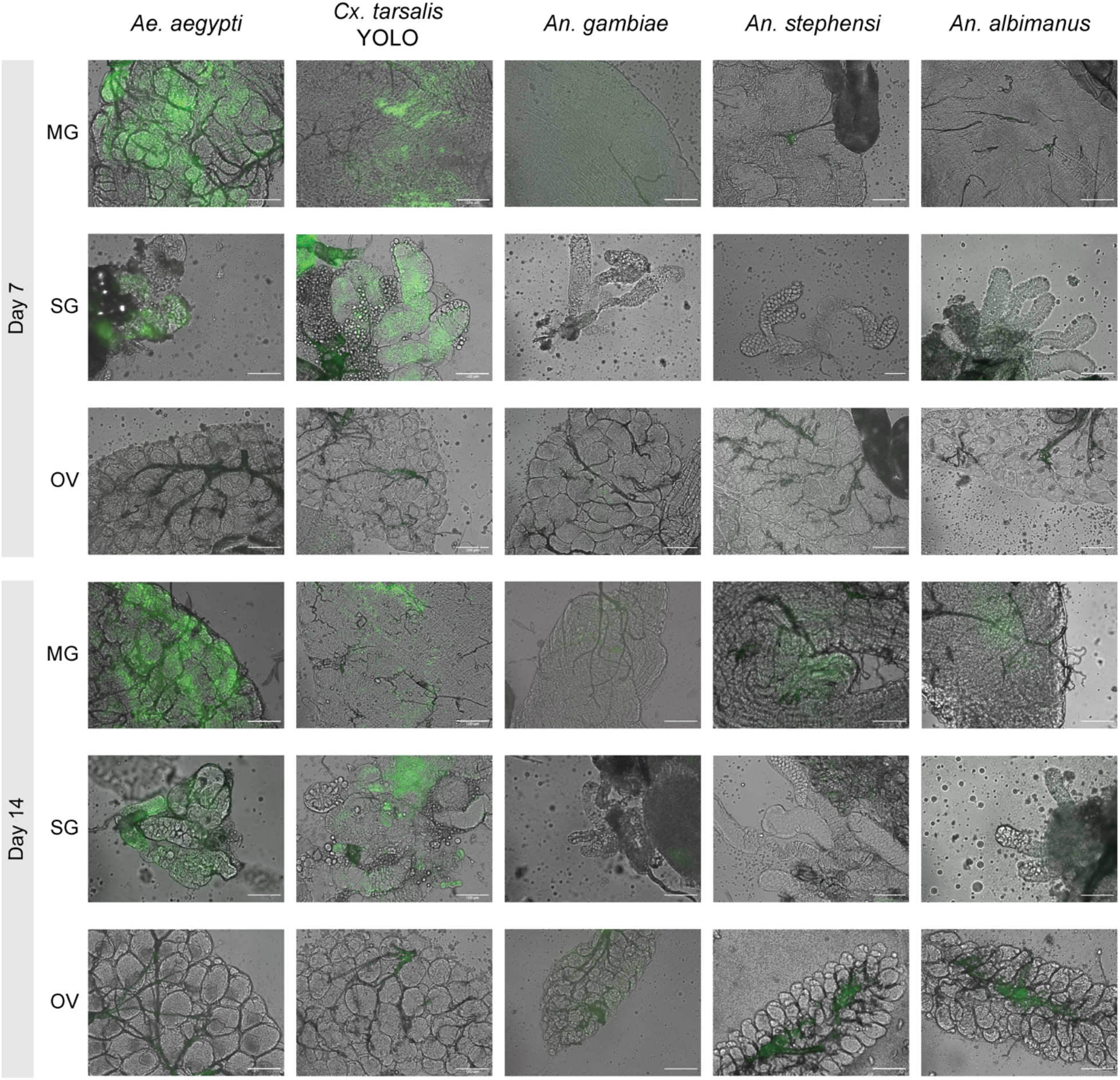
EILV tissue tropism at 7 and 14 dpi in IT injected mosquitoes. Representative images show eGFP fluorescence (or its absence) in midguts (MG), salivary glands (SG) and ovaries (OV) of mosquitoes injected with EILV-eGFP (10^7^ FFU/ml). The brightfield and FITC images have been merged. All scale bars equal 100 μm.

### EILV is not transmitted venereally or vertically in *Cx. tarsalis*

Given the ability of EILV to infect ovaries of *Cx. tarsalis* (YOLO) post-oral infection, we next evaluated whether this virus could be transmitted vertically from females to offspring, and also if EILV could be spread from females to males during mating. Parental *Cx. tarsalis* female mosquitoes (n=15) were orally challenged with EILV-eGFP and then later fed two non-infectious bloodmeals to trigger gonotrophic cycles. After their final oviposition, parental females were dissected and screened for EILV by examining midgut tissue for eGFP fluorescence. We found 80% of parental females remained infected at the midgut level (Table 4). Initially uninfected parental males (n=45) were harvested post-mating and screened for EILV by fluorescence microscopy, FFA, and RT-PCR, and none were found to harbor EILV, indicating no venereal transmission occurred in our assays (Table 4 and Fig. S2B). We next searched for instances of vertical transmission among F1 offspring of infected females by microscopy, FFA, and RT-PCR. The midguts of adult female F1s that emerged from egg raft 2 (ER2; n=55) and ER3 (n=35) were screened for EILV by examining them for eGFP fluorescence. No eGFP was detected in any of the screened F1 midguts, suggesting no vertical transmission occurred (Fig. S2A). FFAs were performed on pooled homogenized female F1 mosquito samples, but no FFUs were detected (Table 4). Similarly, pooled samples from ER2 and ER3 were also negative for EILV RNA by RT-PCR (Fig. S2B) further confirming that EILV was not transmitted vertically.

**TABLE 4.**
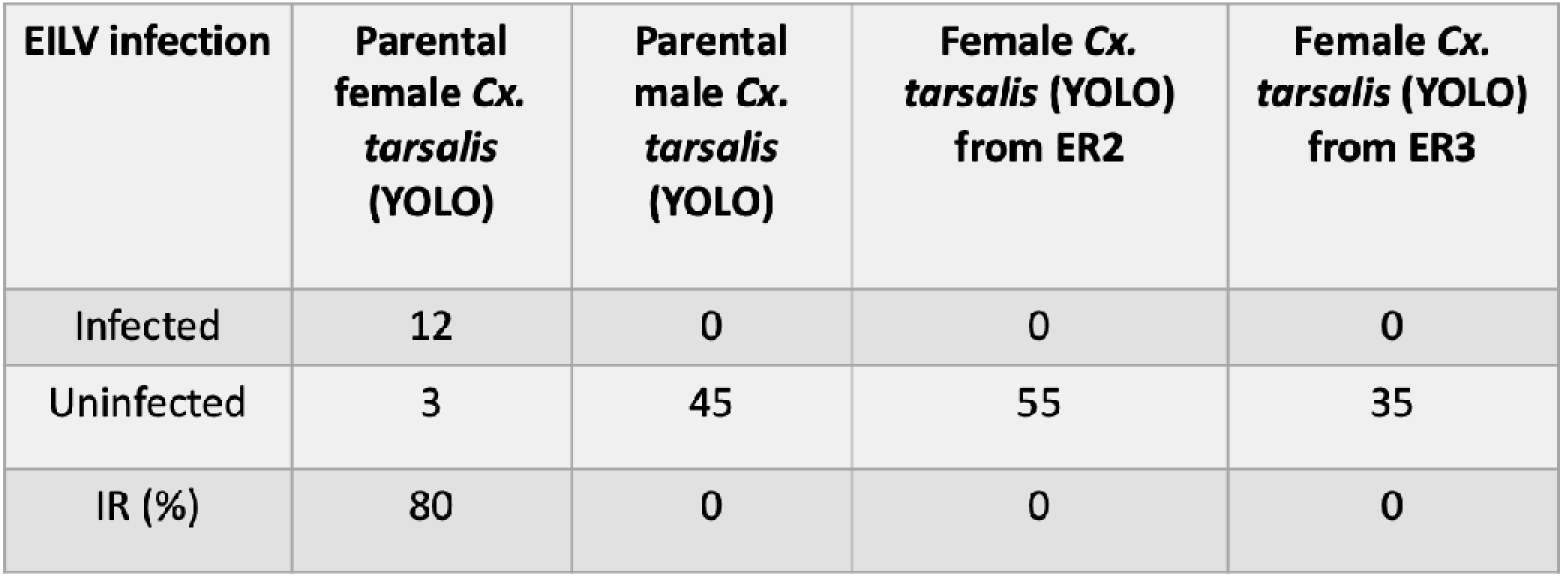
EILV presence in parental and F1 *Cx. tarsalis* (YOLO) to detect vertical and venereal transmission

| **EILV infection** | **Parental female *Cx. tarsalis* (YOLO)** | **Parental male *Cx. tarsalis* (YOLO)** | **Female *Cx. tarsalis* (YOLO) from ER2** | **Female *Cx. tarsalis* (YOLO) from ER3** |
| --- | --- | --- | --- | --- |
| Infected | 12 | 0 | 0 | 0 |
| Uninfected | 3 | 45 | 55 | 35 |
| IR (%) | 80 | 0 | 0 | 0 |

### Reptile cell lines are refractory to EILV infection

Mammalian, avian, and amphibian cell lines were previously shown to be refractory to EILV infection, but reptile cell lines have not been investigated. As mosquitoes can and do feed on reptiles, we investigated the hypothesis that reptiles may be a vertebrate host for EILV. We attempted to infect two reptile cell lines – TH-1, Subline B1 (derived from the box turtle *Terrapene carolina*) and VH 2 cells (derived from Russel’s viper *Daboia russelii*) with EILV-eGFP (10^7^ FFU/ml) and screened for EILV infection at 24, 72 and 120 hrs post-infection under a fluorescence microscope. No eGFP expression was noted in either cell line suggesting that reptile cells are not competent for EILV infection and/or replication.

### *Manduca sexta* caterpillars are not susceptible to EILV infection

Because EILV was found in mosquito saliva and because mosquitoes sometimes bite *M. sexta* caterpillars, we tested *Manduca sexta* caterpillars for EILV host competence. Caterpillars were injected with EILV-eGFP (n=10) or 1xPBS (n=10) and screened for EILV infection at 7 dpi by examining abdominal tissue near the injection site under a fluorescence microscope. No eGFP expression was noted in the injected caterpillars (Fig. S3). Additionally, FFAs were performed on homogenized injected *Manduca sexta* samples and controls, and no FFUs were found. Our results suggest that EILV is unable to infect the invertebrate *Manduca sexta*.

## DISCUSSION

EILV, an insect-specific alphavirus unable infect vertebrates, has the potential to be used as a tool against pathogenic viruses in mosquito vectors^3,7,19^. The goal of this study was to better understand the host range and transmission route(s) of EILV. Here we report that anophelines *An. gambiae, An. albimanus* and *An. stephensi* were not susceptible to EILV by oral infection but were susceptible when the virus was injected intrathoracically. *Ae. aegypti*, on the other hand, was susceptible to EILV by both oral and IT infection routes in our study. These results comport with the findings of a previous study on *An. gambiae* and *Ae. aegypti*^8^. In a new finding, we identify *Cx. tarsalis* as a competent host for EILV.

*Culex tarsalis* (YOLO) was the most competent host for EILV in our study, irrespective of infection route. When challenged orally or IT, *Cx. tarsalis* showed higher infection prevalence compared to other species, and it also frequently presented with higher viral loads (i.e., viral titer). *Culex tarsalis* was also the only mosquito species in our study to transmit EILV via saliva, where the virus was found post-oral and post-IT infection. Only a handful of other insect-specific alphaviruses have been found, and aside from EILV, each was isolated from within the genus *Culex*^4–6^. Our results are consistent with an emerging picture of insect-specific alphaviruses being primarily associated with *Culex*, as is a previous report of EILV infections in *Cx. pipiens* in the wild^12^. However, *Cx. quinquefasciatus*, a closely related species^20^, has low competence for EILV^8^.

Although insect-specific viruses are thought to be adapted to a single host^11,12,21–25^, it remains possible that EILV does not match this framework, instead infecting multiple mosquito taxa. While most ISVs are flaviviruses^26^, only a few insect-specific alphaviruses have ever been described^4,6^, and they may not conform to the patterns described in other ISVs. Mosquito species reported to have host competence for EILV now include An. *coustani*^3^, *Cx. pipiens*^12^, *Cx. tarsalis*, and *Ae. aegypti*^8^, and horizontal transmission through invertebrate hosts or via shared habitats could potentially sustain a multi-host ISV. However, further work is needed to determine the mosquito host use of EILV in nature.

We found that susceptibility of *Cx. tarsalis* to EILV was shared by diverse genetic strains of this species. Specifically, KNWR, another laboratory adapted *Cx. tarsalis* strain, and wild *Cx. tarsalis* from Yolo County, CA, USA (lab reared from egg and larvae stage for one life cycle) both became infected following oral exposure to EILV. However, there were differences among the three tested *Cx. tarsalis* strains. For example, the EILV DIRs of KNWR and wild *Cx. tarsalis* were significantly lower than those of YOLO at 14 days post-oral infection. Congruent to our findings, the susceptibility of mosquitoes to alphaviruses by oral infection is known to vary between strains of the same mosquito species^27–29^. Most strikingly, wild *Cx. tarsalis* mosquitoes had a significantly lower IRs across time, with EILV present in the saliva of wild mosquitoes less often, even when they did become infected. One possible explanation is that the laboratory-adapted colonies have less genetic variation than their wild counterparts, increasing their susceptibility to viruses^27,30–32^.

Our characterization of tissue tropism implies the presence of barriers to EILV infection that differ among species. We find anophelines *An. gambiae, An. albimanus* and *An. stephensi* were not susceptible to EILV by oral infection but were susceptible when the virus was injected into the thorax, indicating the presence of a midgut infection barrier against EILV in those species. In *Ae. aegypti, we* did not detect EILV in saliva despite their salivary gland tissue being infected, suggesting a salivary gland escape barrier against EILV ^33^. Overall, these findings agree with previous work^9^, though more studies are needed verify these barriers.

Interestingly, we observed the presence of EILV in the ovaries of IT-infected *An. gambiae* at 7 and 14 dpi—a tissue tropism not observed in the previous EILV host range study^8^. This difference in ovarian susceptibility could stem from the different *An. gambiae* strains that were tested^8^. Our study used the Keele strain, which was developed by the balanced interbreeding of four An. *gambiae sensu stricto* strains, one of which is the G3 strain used in the previous study^34^. Thus, there was genetic overlap in the strains tested. However, the Keele strain has significantly higher allelic diversity, which could underlie the observed disparity in EILV tissue tropism^35,36^.

Because insect-specific viruses are believed to be adapted to a single host, they are thought to depend on vertical transmission (and, to a lesser extent, venereal transmission) to maintain themselves in host populations^11,12,21–25^. Moreover, the efficiency of vertical transmission of insect-specific viruses seems to be higher^21^ than viruses that spread by horizontal transmission ^37–39^. For example, an insect-specific flavivirus in *Cx. pipiens* (CxFV) was found to vertically transmit at 100% efficiency via transovarial transmission (TOT)^21^. Consistent with the hypothesis that it transmits vertically, EILV has been detected in *Cx. pipiens* larvae in the wild^14^. Moreover, the presence of EILV in the ovaries of *Cx. tarsalis* (YOLO) and *Cx. tarsalis* (KNWR) in the present study is preliminary evidence of TOT, though the virus was not found in the ovaries of wild *Cx. tarsalis* post-oral infection. This absence could indicate a barrier to EILV infection similar to the midgut escape barrier—though one that has been lost in the tested laboratory-adapted strains^40^. Further experiments are needed to verify and characterize this variation in ovarian tropism, and to determine if EILV may be transmitted vertically under different conditions than those tested here.

Strikingly, no EILV was detected in F_1_ progeny of infected *Cx. tarsalis* females in our vertical transmission study. Moreover, no venereal transmission from infected females to their male partners was noted. This may indicate that *Cx. tarsalis* (YOLO) is not a native host for EILV and vertical transmission is restricted to a more specialized (unknown) host. Another possibility is that persistently EILV-infected populations may be more efficient at vertical transmission due to physiological changes caused by persistent natural infection. There is some empirical evidence for the latter hypothesis: CxFV (an ISV) was transmitted vertically by naturally infected *Cx. pipiens* but not by IT-infected naïve *Cx. pipiens*^21^.

In *Cx. tarsalis* (YOLO), EILV tissue tropism did not differ between oral and IT infection routes, with the exception that no EILV was found in IT-infected *Cx. tarsalis* (YOLO) ovaries. This suggests that blood-feeding is necessary for EILV to infect the ovaries. Mosquito ovaries undergo oogenesis post-blood feeding, during which the ovaries expand as nutrients enter to form the yolk in a process called vitellogenesis^41^. These morphological changes may be used by EILV to infect the ovaries. Similarly, plant viruses are known to use vitellogenesis to infect the ovaries of white flies, and in mosquitoes, protein and DNA cargo have been transported into the ovaries using vitellogenin^42,43^. However, more research is required to determine the mechanism of infection of the ovaries by EILV in *Cx. tarsalis* (YOLO).

EILV is thought to have evolved into a single-host virus from a dual-host ancestor, like the insect-specific flaviviruses that cluster with dual-host viruses^3,7,44^. Phylogenetically, EILV is most closely related to alphaviruses that infect both mosquitoes and vertebrates^3^, but EILV lost its ability to infect vertebrates. The presence of EILV in the saliva of *Cx. tarsalis* (YOLO), *Cx. tarsalis* (KNWR), and wild *Cx. tarsalis* may be a remnant of its past ability to transmit from mosquitoes to vertebrates horizontally^3,7,8^. Alternatively, EILV may still be a dual- or multi-host virus, horizontally transmitted between mosquitoes and another vertebrate or invertebrate host. Our data suggests that reptile cells are not competent for EILV infection, suggesting that reptiles are not a vertebrate host for the virus. Recent research has shown that mosquitoes not only feed on vertebrates but can also feed on invertebrates such as worms, leeches, and caterpillars^15,45^. These diverse food sources increase the possible host ranges of arboviruses. While we find that EILV infection is not supported by *Manduca sexta* larvae, another invertebrate may be susceptible to EILV. Further research could explore the role of diverse invertebrates in the maintenance of arboviruses in nature.

Our study adds to an emerging picture that EILV may have non-standard characteristics for an insect-specific virus. Ostensibly, it can infect multiple mosquito species, including in the genera *Anopheles*^9^, *Aedes*^9^, and—as we demonstrate here—Culex^12^. However, its transmission route(s) remain elusive. There is limited evidence for transovarial transmission, the dominant route for better-characterized ISVs, but EILV was also found in salivary glands and was secreted in saliva—revealing the potential for horizontal transmission. Notably, we find EILV replicating in *Cx. tarsalis* ovary tissue, the first report of EILV in the ovary of any species. Because it efficiently infects *Culex tarsalis—and* grew in all examined tissues—EILV may prove a useful tool to combat pathogenic viruses transmitted by this mosquito, such as West Nile Virus and Western Equine Encephalitis.

## MATERIALS AND METHODS

### Cells and cell culture

The An. *albopictus* mosquito cell line C6/36 was propagated at 28°C with no CO_2_ in complete RPMI medium, which comprised Roswell Park Memorial Institute 1640 (RPMI 1640) medium (Gibco/Thermo Fisher Scientific, Waltham, MA, USA) supplemented with 10% (v/v) fetal bovine serum (FBS) (Gibco/Thermo Fisher Scientific), penicillin (100 U/ml) (Gibco/Thermo Fisher Scientific), streptomycin (100 μg/ml) (Gibco/Thermo Fisher Scientific), and 2% (v/v) tryptose phosphate broth (Sigma-Aldrich).

TH-1, Subline B1 (*Terrapene carolina*) and VH 2 (*Daboia russellii*) cells were acquired from ATCC and propagated in complete Minimum Essential Media (MEM, (Gibco/Thermo Fisher Scientific) supplemented with 10% (v/v) fetal bovine serum (FBS) (Gibco/Thermo Fisher Scientific), penicillin (100 U/ml) (Gibco/Thermo Fisher Scientific), streptomycin (100 μg/ml) (Gibco/Thermo Fisher Scientific). TH-1, Subline B1 cells were maintained at 28°C with no CO_2_ while VH 2 cells were maintained at 30°C with 5% CO_2_.

### Viral cDNA clone and virus rescue

We used an EILV (strain EO329) cDNA clone with an enhanced green fluorescent protein inserted in the hypervariable region of nsp3 of the EILV genome (EILV-eGFP) for all experiments. The EILV-eGFP cDNA clone was obtained from the World Reference Center for Emerging Viruses and Arboviruses at the University of Texas Medical Branch.

EILV-eGFP was rescued as previously described with some modifications^7^. The cDNA clone (10 μg) was linearized using NotI (New England BioLabs [NEB], Ipswich, MA, USA) then purified and concentrated using the DNA Clean and Concentrator-25 kit (Zymo Research, Irvine, CA, USA). The linearized cDNA was transcribed using MEGAscript SP6 Transcription Kit (Invitrogen/Thermo Fisher Scientific) with the addition of an m7G(5⍰)ppp(5⍰])G RNA cap (NEB). Transcription was carried out at 42°C for 2 hours (h) on a thermocycler. The transcribed RNA was then purified using a MEGAclear Transcription Clean-Up Kit (Invitrogen/Thermo Fisher Scientific) and stored at −80 °C.

Before transfection, C6/36 cells were seeded in a T75 flask at a density of 6 × 10^7^ cells and incubated overnight at 28°C with no CO_2_ to achieve ~70-80% confluency. Transfection of EILV-eGFP RNA was performed using the TranIT-mRNA Transfection Kit (Mirus Bio, Madison, WI, USA). We verified EILV-eGFP infection in transfected cells by examining them for eGFP fluorescence using a Zeiss Axiovert S 100 equipped with a Fourier-transform infrared spectroscopy (FTIR) filter. To collect infectious virus, cell supernatant was harvested 5 days post-transfection and centrifuged for 10 min at 3000 x *g* at 4°C in a swinging bucket rotor centrifuge to remove cell debris. The supernatant was then aliquoted and stored at −80°C. EILV-eGFP viral titer was quantified by focus-forming assays (described below).

To infect reptile cell lines, 12 well tissue culture plates were seeded with 1X10^7^ TH-1, Subline B1 or VH 2 cells per well and incubated overnight at 28°C with no CO_2_ and 30°C with 5% CO_2_ respectively to achieve ~70-80% confluency. The reptile cells were washed with MEM containing no FBS and inoculated with 0.5 ml of EILV-eGFP (10^7^ FFU/ml) and incubated at their respective growth conditions for 1h. Then, the virus was removed, and replaced with complete MEM. We scored EILV-eGFP infection in the reptile cells by examining them for fluorescence at 24, 72, 120 hrs post-infection using a Zeiss Axiovert S 100 equipped with a Fourier-transform infrared spectroscopy (FTIR) filter.

### Mosquitoes and mosquito rearing

We examined competence for EILV infection in five mosquito species: *Aedes aegypti, Culex tarsalis, Anopheles gambiae, An. stephensi*, and *An. albimanus*. A positive control species for EILV infection^8^, *Ae. aegypti* (Rockefeller strain), was provided by Johns Hopkins University (Baltimore, MD, USA). *Anopheles gambiae* (Keele strain) was obtained from The National Institutes of Health (Bethesda, MD, USA) and served as a negative control^8^. *Anopheles albimanus* (STELCA strain) was obtained from BEI Resources (Manassas, VA, USA). *Anopheles stephensi* (Liston strain) was acquired from Johns Hopkins University (Baltimore, MD, USA). We tested three strains of *Cx. tarsalis:* YOLO strain (from BEI Resources, Manassas, VA, USA), KNWR strain (from Christopher Barker’s laboratory, UC Davis School of Veterinary Medicine, Davis, CA, USA) and wild *Culex tarsalis* (from Sacramento-YOLO mosquito and vector control district, Elk Grove, CA, USA).

All mosquitoes were reared and maintained at the Millennium Sciences Complex (The Pennsylvania State University, University Park, PA, USA). *Culex tarsalis* (YOLO and KNWR strain) larvae were reared in 30×30×30 cm cages and maintained at 25°C ± 1°C, 16:8 h light: dark diurnal cycle with 80% relative humidity. The other mosquitoes were reared in 30×30×30 cm cages in a walk-in environmental chamber maintained at 27°C ± 1°C, 12:12 h light: dark diurnal cycle with 80% relative humidity. Wild *Culex tarsalis eggs* collected in the field were reared for one generation in the laboratory. *Aedes aegypti, An. albimanus*, and *Cx. tarsalis* (YOLO and KNWR strain) larvae were fed Koi pellets (Tetra, Melle, Germany). *Anopheles gambiae and Cx. tarsalis* (wild) larvae were fed TetraMin (Tetra, Melle, Germany). A slurry made with Tetramin and baker’s yeast (1:2 by volume) was used to feed *An. stephensi* larvae. All adult mosquitoes were provided with 10% sucrose solution on cotton balls *ad libitum*.

### Mosquito infections

For both infection routes, each mosquito species was challenged separately using aliquots of the same virus stock. For each species, we challenged two replicate batches of mosquitoes. Half of the mosquitoes from each batch were then sampled at each of the two time points (i.e., 7 dpi and 14 dpi).

#### Oral infections

Adult female mosquitoes (3–5 days post-emergence) were sugar-starved for 24 h then fed an infectious blood meal comprised of 1:1 anonymous human blood (BioIVT, Westbury, NY, USA) and 10^7^ FFU/ml EILV-eGFP at 37°C using a water-jacketed membrane feeder. Mosquitoes were then cold-anesthetized, and fully engorged mosquitoes were counted and placed in cardboard cup cages until processing.

#### Intrathoracic infections

Adult female mosquitoes (3–5 days post-emergence) were cold-anesthetized and placed on a glass slide on top of a chill block maintained at 4°C. Under an Olympus SZX7 Stereo Microscope, each mosquito was then injected in the thorax with 100 nl of EILV-eGFP virus (107 FFU/ml) at a rate of 100 nl/s using a Nanoject III (Drummond Scientific Company, Broomall, PA, USA). Injected mosquitoes were counted and placed in a sealed cardboard cup cage until processed.

### Imaging

Mosquitoes challenged with EILV-eGFP were dissected on days 7 and 14 dpi. For each time point, midgut, ovaries, and salivary glands of each mosquito were dissected out into 50 μl of 1xPBS on a glass slide. A cover slip was placed over the mosquito tissues for imaging. EILV infection status in organs was determined by examining tissue for eGFP fluorescence using an Olympus BX41 inverted microscope equipped with a FITC filter. The organs were also imaged under brightfield illumination.

### Host competence assays

The host competence of each mosquito species for EILV at 7 and 14 dpi was determined as previously described^16–18^. Briefly, challenged mosquitoes were anesthetized using triethylamine (Sigma-Aldrich, St. Louis, MO, USA) and forced to salivate into a capillary glass tube containing FBS and 50% sucrose mixed at a ratio of 1:1 for 30 min. After 30 min, the saliva was pushed into a 2 ml microcentrifuge tube containing 100 μl mosquito dilutant (1xPBS mixed with 20% of FBS, 100 μg/ml of streptomycin, 100 units/ml of penicillin, 50 μg/ml gentamicin [Gibco/Thermo Fisher Scientific], and 2.5 μg/ml Amphotericin B [Gibco/Thermo Fisher Scientific]). Next, bodies and legs were collected and placed separately into 2 ml microcentrifuge tubes, each containing 300 μl mosquito dilutant and a 4.5 mm zinc-plated steel bead (Daisy Outdoor Products, Rogers, AR, USA). Samples were briefly stored on ice, then leg and body samples were homogenized using a TissueLyser II (Qiagen, Hilden, Germany) at 30 Hz for 2 min. Homogenized samples were centrifuged at 6000x *g* at 4°C for 5 min using a benchtop microcentrifuge. All samples were stored at −80°C until quantified by FFA (described below). From FFA counts, the IR was calculated as the proportion of infected mosquitoes among the total number of engorged mosquitoes, the DIR the proportion of infected mosquitoes with virus-positive legs, the TR the proportion of mosquitoes with virus-positive saliva among those with virus-positive legs, and the TE the proportion of mosquitoes with virus-positive saliva among the total number of mosquitoes engorged.

### Focus-forming assays (FFA)

We quantified EILV-eGFP titers in samples using focus-forming assays. C6/36 cells were seeded in 96-well plates at a density of 1 × 10^5^ cells/well and incubated at 28°C with no CO_2_ overnight. Complete RPMI media was then removed, and serially diluted (10^-1^ to 10^-4^) mosquito samples in serum-free RPMI media were then added (30 uL) in duplicate to the prepared cells. Saliva samples were not diluted. The cells were incubated at 28°C with no CO_2_ for 1 h. The samples were then removed, and cells were covered with 100 μl of RPMI containing 0.8% methylcellulose (Sigma-Aldrich). The infected cells were incubated at 28°C without CO_2_ for 48 h. The infected C6/36 cells were fixed using 50 μl of 4% formaldehyde (Sigma-Aldrich) in 1xPBS for 30 min at room temperature. Fixed cells were washed two times with 100 μl of 1xPBS, and finally, 50 μl of 1xPBS was added to the cells (to prevent drying) and fluorescent (i.e., EILV-eGFP) foci were counted using a FITC equipped Olympus BX41 inverted microscope.

### Vertical transmission assays

Vertical transmission assays were carried out as previously described^17^ with some modifications. We collected eggs from virus-challenged females at three time points to evaluate vertical transmission of EILV. Adult female *Cx. tarsalis* (3–5 days post emergence) were fed an artificial infectious blood meal supplemented with EILV-eGFP (10^7^ FFU/ml) to induce EILV infection. Blood-fed mosquitoes were cold-anesthetized and fully engorged females were counted and sorted into sealed cardboard cups. EILV-eGFP fed females were then fed non-infectious blood meals at both 7 and 14 dpi. Oviposition containers, consisting of a wide-mouth plastic cup filled halfway with DI water, were placed inside the cages 4 days after each feed and were collected 5 days after each feed (i.e., mosquitoes had ~24 h of access). The egg rafts from the three gonotrophic cycles were labeled as ER1, ER2, and ER3. ER1 was discarded while ER2 and ER3 were hatched and reared to the adult stage. Midguts were dissected from parental female and male mosquitoes as well as from adult offspring (3 days post-emergence) reared from ER2 and ER3 eggs. Midguts were evaluated for EILV-eGFP infection by fluorescence microscopy as described above. The carcasses were then collected in pools of five in a 2 ml microcentrifuge tube containing 500 μl of mosquito diluent and one 4.5 mm zinc-plated steel bead. Carcass samples of parental males and offspring included midguts, which were returned post imaging. The samples were homogenized using a TissueLyser II (Qiagen) at 30 Hz for 2 min then centrifuged at 6000x *g* at 4°C for 5 min using a benchtop microcentrifuge. All samples were quantified by both FFA, as described above, as well as by RT-PCR.

### RT-PCR to detect EILV infection

RNA was extracted from test and control mosquito samples using the Direct-zol RNA Miniprep Kit (Zymo Research). The positive and negative controls for the RT-PCR were RNA extracted from an FFA-confirmed EILV-eGFP positive mosquito and an FFA-confirmed EILV-eGFP negative mosquito (EILV-eGFP challenged), respectively. The extracted RNA samples were quantified using a NanoDrop ND-1000 (NanoDrop Technologies/Thermo Fisher Scientific), then the template was amplified using the OneStep RT-PCR Kit (Qiagen) with the EILV-specific primers 5’-CGA CGA TGA CCG GAG AAG AG *-3’* and reverse primer 5’-AAG ACT CGG TCT GCC TGC −3’. Amplicons were analyzed using gel electrophoresis.

### *Manduca sexta* rearing and infection

Early-stage (L2-L3) *Manduca sexta* larvae were acquired from Rudolf Schilder at Pennsylvania State University (University Park, PA, USA). The larvae were reared at room temperature in 30×30×30 cm plastic containers with air holes and fed an artificial diet (Frontier Agricultural Sciences, Newark, DE, USA) *ad libitum*.

Larvae were anesthetized on ice and the injection site was swabbed with 70% ethanol. Larvae were then injected with either 10 μL of 1×10^7^ FFU/ml EILV-eGFP or 1x phosphate-buffered saline (PBS, Gibco/Thermo Fisher Scientific, negative control) between the abdominal prolegs using sterile insulin syringes. Injected treatment and control *Manduca sexta* larvae were sacrificed 7 days post injection at −80°C for 10 min. We initially screened larvae for EILV infection by dissecting the tissue near the injection site and examining it for eGFP fluorescence (indicating expression of EILV-eGFP) using an Olympus BX41 inverted microscope equipped with a FITC filter. Post imaging, samples were homogenized using a motorized homogenizer and pestle, followed by centrifugation for 10 min at 3000 x *g* at 4°C in a swinging bucket rotor centrifuge. All samples were stored at −80°C until processed. EILV-eGFP viral titers were measured by FFA as described above.

### Statistical analysis

Fisher’s exact tests were used to evaluate differences in the infection rate (IR), dissemination rate (DIR), transmission rate (TR), and transmission efficiency (TE) between mosquito species orally and IT infected with EILV-eGFP. The EILV viral titers of body, leg, and saliva samples were compared using Mann-Whitney U tests. All statistical tests were run using GraphPad Prism version 9.0.4.

## ACKNOWLEDGEMENTS

The authors acknowledge Mrs. Amelia Romo for her assistance with mosquito rearing, and Dr. Sarah Wheeler from Sacramento-YOLO mosquito and vector control district, Elk Grove, CA, USA, for providing us with *Cx. tarsalis* eggs from wild-caught gravid female *Cx. tarsalis* mosquitoes, and Dr. Rudolf Schilder at Pennsylvania State University (University Park, PA, USA) for supplying us with *Manduca sexta* larvae and technical assistance rearing them. This study was supported by NIH/NIAID grants R01A1128201 and R01AI150251, NSF/BIO grant 1645331, USDA Hatch Project 4769, a grant with the Pennsylvania Department of Health using Tobacco Settlement Funds, and funds from the Dorothy Foehr Huck and J. Lloyd Huck endowment to JLR.

## ABBREVIATIONS

CxFV: Culex flavivirus
DIR: dissemination rate
dpi: days post infection
eGFP: enhanced green fluorescent protein
EILV: Eilat virus
FFA: focus forming assay
FFU: focus forming unit
ISV: insect specific virus
IR: infection rate
IT: intrathoracic
MG: midgut
OV: ovary
SG: salivary gland
TE: transmission efficiency
TOT: transovarial transmission
TR: transmission rate

