## Supplementary material for "*Culex tarsalis* is a competent host of the insect-specific alphavirus Eilat virus (EILV)": Sup figures

### Mosquito host-range and transmission routes of Eilat virus

<u>This PDF contains:</u>

Supplementary Figures S1-S3

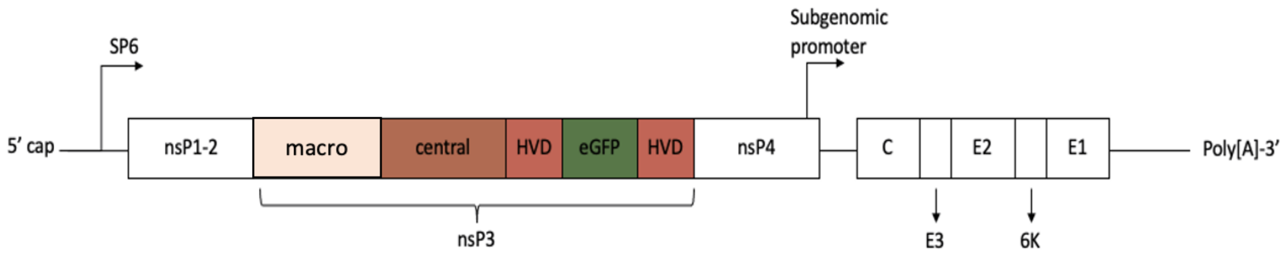


**FIG S1** A schematic diagram of the EILV-eGFP cDNA clone used in all infection experiments. The insertion of the eGFP coding sequence into HVD of nsP3 of the EILV genome (strain O329) is illustrated. This template was used for the rescue of EILV-eGFP and was obtained from the UTMB World Reference Center for Emerging Viruses and Arboviruses.

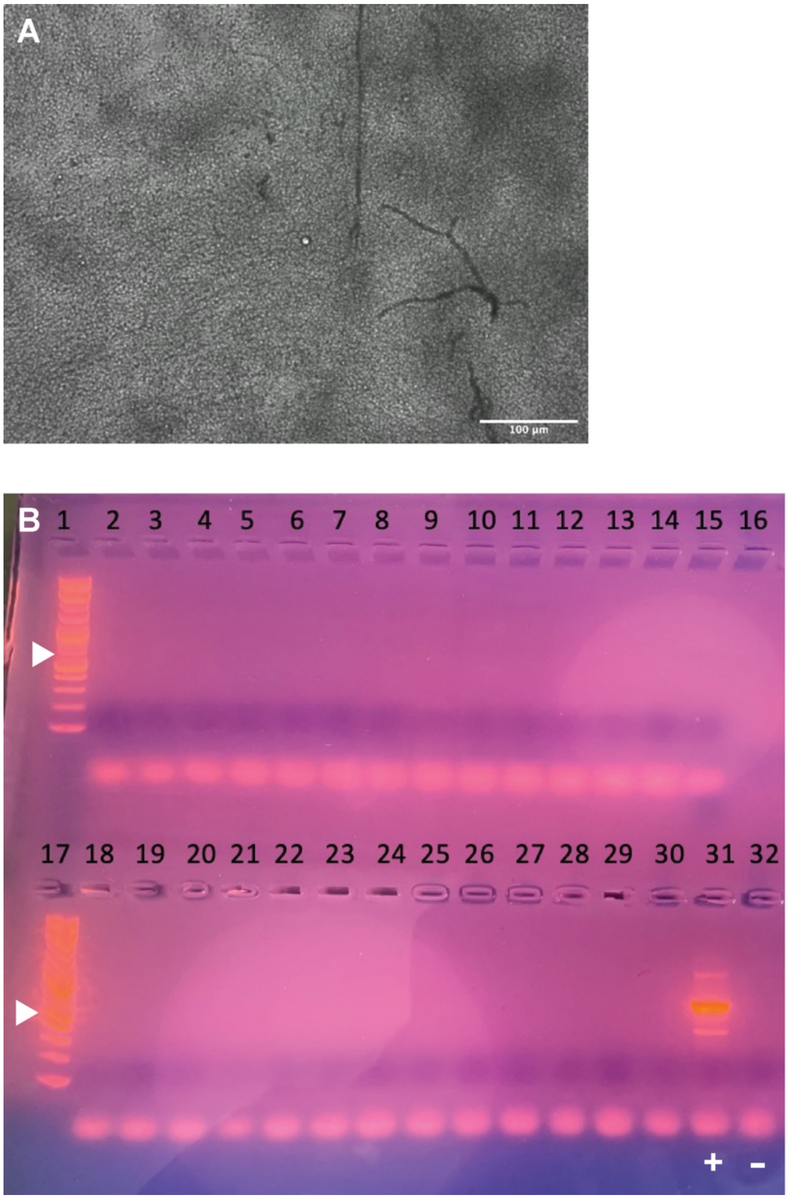


**FIG S2** Additional data related to vertical and venereal transmission in *Cx. tarsalis* (YOLO).
A) Merged fluorescence and brightfield images of the midgut of a female mosquito emerged from ER3. The scale bar equals 100 µm.
B) Gel electrophoresis of EILV RT-PCR products from pooled ER2, ER3, and paternal male samples. Wells 1 and 17 are 1 kb plus ladder, where white arrowheads mark the position of the 600 bp band. Wells 2-12 are pooled samples from ER2. Wells 13-15 and 18-21 are pooled samples from ER3. The pooled paternal male samples are present in wells 22-30. The 605 bp positive control (marked '+') in well 31 is amplified from EILV-positive mosquito samples. The negative control (marked '-') in well 32 used an EILV-negative mosquito sample as template.

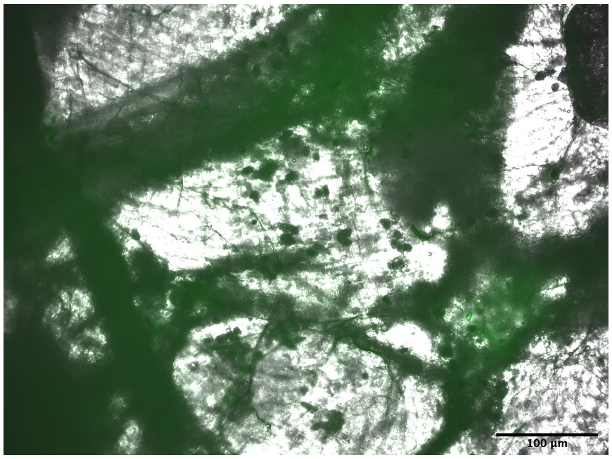


**FIG S3** Merged fluorescence and brightfield images of a *Manduca sexta* tissue section at 7 dpi dissected from near the site of EILV-eGFP ($10^7$ FFU/ml) injection. The scale bar equals 100 µm. Green signal is general autofluorescence.
